# A new component of the DNA damage response biofilm axis is a TetR-like DNA damage response regulator in *Acinetobacter baumannii*

**DOI:** 10.1101/2025.06.05.658052

**Authors:** Brian Nguyen, Mark W. Soo, Guillermo Antunez Tierney, Elizabeth A. Amorelli, Austen L. Herlihy, Veronica G. Godoy

## Abstract

*Acinetobacter baumannii* is an opportunistic pathogen that employs two main strategies to evade antibiotic treatment: developing antibiotic resistant through the DNA damage response (DDR), and forming biofilms, which are protective bacterial multicellular communities. Previously, we demonstrated that RecA, a key regulator of the DDR, connects the DDR and biofilm formation, with RecA levels inversely correlated with biofilm formation. In this study, we identify another DDR regulator, EppR— a recently characterized TetR-family transcriptional repressor— as also playing a role in biofilm formation. We show that an *eppR*-deficient strain is unable to form biofilms due to reduced expression of genes encoding adhesive pili. This occurs because EppR influences intracellular RecA levels. Our findings provide further insight into RecA regulation and the link between the DDR and biofilms.

## Introduction

*Acinetobacter baumannii* is a gram-negative nosocomial ESKAPE pathogen (Boucher et al., 2009). These pathogens are known for their ability to become multidrug resistant, leading to infections that are difficult to treat, therefore causing great strain on healthcare systems. *A. baumannii* can avoid eradication by antibiotics through a variety of measures, including low membrane permeability to antimicrobials (Bou et al., 2000; Clark, 1996; Fernandez-Cuenca, 2003; Vila et al., 2007), efflux pumps (Pérez-Varela et al., 2018; Richmond et al., 2016; Roca et al., 2009; Xu et al., 2019), and production of various classes of β-lactamases (Raible et al., 2017). Additionally, previous work has shown that in *A. baumannii* (Norton et al., 2013) and other bacteria there is a causal relationship between the DNA damage response (DDR) and antibiotic resistance acquisition through mutagenesis of antibiotic targets due to the activity of error-prone DNA polymerases in response to DNA damage (Boshoff et al., 2003; Cirz et al., 2007; Cirz, O’Neill, et al., 2006).

*A. baumannii* also resists antibiotic challenge and survives desiccation through the formation of multicellular communities or biofilms (Amin et al., 2019; Bhattacharyya et al., 2013; Li et al., 2021) by adhering to various biotic and abiotic surfaces present in hospital settings (Ahmad et al., 2023; Ronish et al., 2019; Tomaras et al., 2003). While some of the key biofilm genes are known, the mechanisms underlying the biofilm lifecycle are still not fully understood. Previous studies have shown that cell attachment is facilitated by an archaic chaperon-usher pili encoded in a single operon, Csu, encoding the proteins CsuAB, A, B, C, D, and E (A1S_2218 – A1S_2213, ACX60_RS06480 – ACX60_RS06505) (Ahmad et al., 2023; Pakharukova et al., 2018; Tomaras et al., 2003). It is known that the *csu* operon is somehow regulated by the two-component histidine kinase system BfmR/S (A1S_0748 and A1S_0749, ACX_RS14635 and ACX_RS14640); BfmR is required for expression of the *csu* operon (Gaddy & Actis, 2009; Tomaras et al., 2008). The intricacies of this regulation remain unknown as BfmR does not bind to the *csu* promoter (Raustad et al., 2025). We showed that VanR, a regulator of vanillin catabolism, binds to the *csu* promoter and represses its expression. Induction of the *csu* genes occurs when vanillate binds to VanR (Brychcy et al., 2024). The interplay between BfmR and VanR is unknown. We have also shown that Lon protease plays a role in biofilm formation as Lon-deficient strains of *A. baumannii* do not form biofilms (Ching et al., 2019). Furthermore, genes encoding Bap, outer membrane proteins, such as OmpA, and efflux pumps are also involved in biofilm formation (Brossard & Campagnari, 2012; Choi et al., 2008; Gaddy et al., 2009; He et al., 2015). Overall, though great strides have been made to understand biofilm development in *A. baumannii*, much remains unknown.

Remarkably, there is also evidence for a link between biofilm development and the DDR (Linares et al., 2006; Takahashi et al., 1995). While there is a direct correlation between DDR and biofilm development in other bacterial species (Costa et al., 2014; Inagaki et al., 2009; Walter et al., 2015), we have shown that in *Bacillus subtilis*, there is an inverse correlation between the two pathways (Gozzi et al., 2017). We observed that, in *B. subtilis*, biofilm matrix genes are downregulated when the DDR is induced by oxidative stress. DDR-induced cells and cells producing matrix genes are mutually exclusive within biofilms. This strategy would allow DDR-induced cells to leave the biofilm and find less stressful environments. Similarly, in *A. baumannii,* RecA, one of the key DDR regulators, influences biofilm development depending on its protein levels (Ching et al., 2024). In response to DNA damage, cells form two distinct subpopulations, in which one expresses *recA* at high and the other at low levels (MacGuire et al., 2014). High RecA would result in DNA repair, mutagenesis, and biofilm dispersal while low RecA would lead to biofilm formation (Brychcy et al., 2024; Ching et al., 2024). These results demonstrate the existence of a connection between the DDR and biofilm formation. *A. baumannii* can use this DDR-biofilm link to deploy novel strategies to deal with environmental stress.

We recently identified a novel TetR-like transcriptional regulator in *A. baumannii*: EppR (<u>e</u>rror-<u>p</u>rone <u>p</u>olymerase regulator; A1S_1669, ACX60_RS09590). We showed that it is a repressor of *umuDAb*, *umuDC*s, and *umuC* genes, which respectively encode a DDR regulator and various copies of error-prone DNA polymerase V (Nguyen et al., 2025). In this study, we show that EppR also influences biofilms as deletion of EppR results in biofilm deficiency. In this strain, we observe, for example, no Csu pili correlated with low expression of *csuAB*, and reduced expression of various genes previously shown to be involved in biofilms, such as *bfmR* and *lon*. Remarkably, our results indicate that EppR influences intracellular levels of RecA leading us to surmise that this may be how EppR affects biofilm formation.

## Results

### Loss of EppR results in lack of biofilm formation

We previously demonstrated that the DDR-biofilms axis in *A. baumannii* (Ching et al., 2024) involves the DDR regulator UmuDAb (A1S_1389, ACX60_RS11175) (Aranda et al., 2013; Hare et al., 2014) among others. Recently, we identified EppR (<u>E</u>rror-<u>p</u>rone <u>p</u>olymerase regulator; A1S_1669, ACX60_RS09590), another DDR regulator that happens to control *umuDAb* expression (Nguyen et al., 2025). Because UmuDAb is important to both DDR and biofilms, we hypothesized that EppR might also be part of the DDR-biofilm connection.

To test this, we performed a biofilm assay in which cells form a thin pellicle at the air - liquid interface with the following strains: wild-type *A. baumannii* ATCC 17978 (WT), Δ*eppR*, and Δ*eppR* complemented with a low-copy plasmid, pNLAC (Luke et al., 2010) containing *eppR* under its own promoter (Nguyen et al., 2025) (Δ*eppR*c; Table 1). To control for the addition of the plasmid in Δ*eppR*c, WT and Δ*eppR* contain the empty pNLAC plasmid.

**Table 1:**
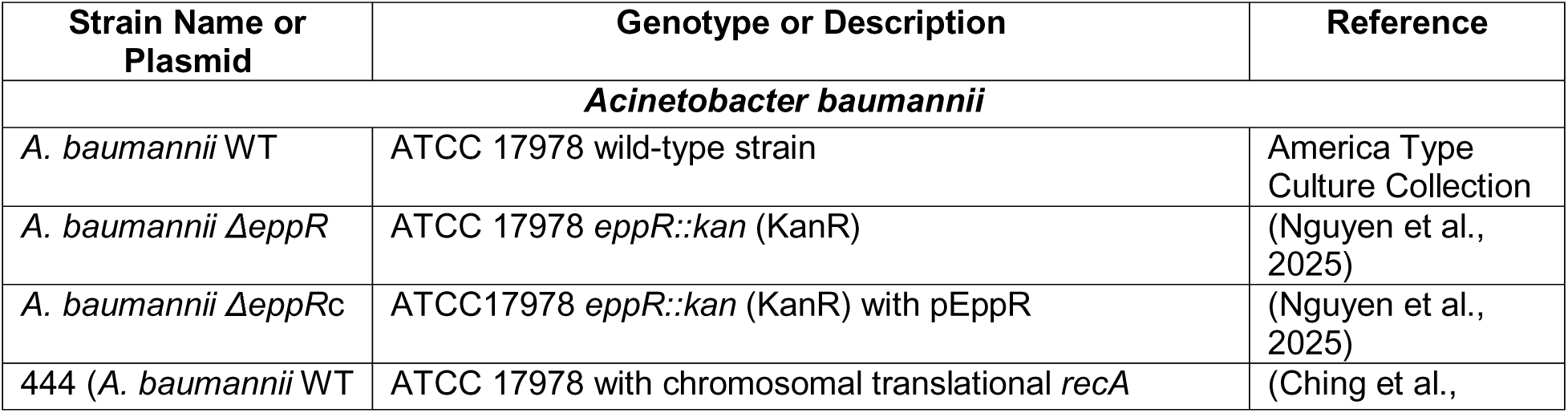
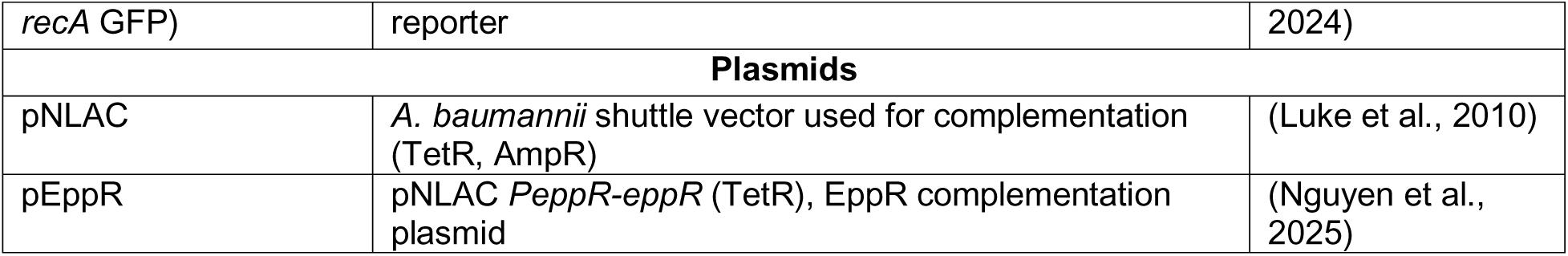
Strain and Plasmid List.

| Strain Name or Plasmid | Genotype or Description | Reference |
| --- | --- | --- |
| *Acinetobacter baumannii* |  |  |
| *A. baumannii* WT | ATCC 17978 wild-type strain | America Type Culture Collection |
| *A. baumannii* ΔeppR | ATCC 17978 *eppR::kan* (KanR) | (Nguyen et al., 2025) |
| *A. baumannii* ΔeppRc | ATCC17978 *eppR::kan* (KanR) with pEppR | (Nguyen et al., 2025) |
| 444 (*A. baumannii* WT | ATCC 17978 with chromosomal translational *recA* | (Ching et al., |
| *recA* GFP) | reporter | 2024) |
| **Plasmids** |  |  |
| pNLAC | *A. baumannii* shuttle vector used for complementation (TetR, AmpR) | (Luke et al., 2010) |
| pEppR | pNLAC *PeppR-eppR* (TetR), EppR complementation plasmid | (Nguyen et al., 2025) |

We found that after 72 hours, WT and Δ*eppR*c formed a noticeable pellicle biofilm, where Δ*eppR*c cells generally formed a smoother looking pellicle. In contrast, in the same time frame Δ*eppR* cells showed no signs of biofilms (Figure 1A). To quantify these biofilms, we performed crystal violet staining to measure cell surface attachment along the sides and bottom of the pellicle biofilms (Ching et al., 2019) (Figure 1B). Δ*eppR* showed a significant 10-fold reduction while Δ*eppR*c showed a modest, but significant, increase of surface-attached cells compared to WT. The lack of biofilm exhibited in Δ*eppR* is not due to a growth deficiency in the strain. Growth experiments show that Δ*eppR* grows similarly to WT (Figure S1). This result agrees with our hypothesis suggesting that EppR is not only involved with the DDR but also influences biofilms.

**Figure 1:**
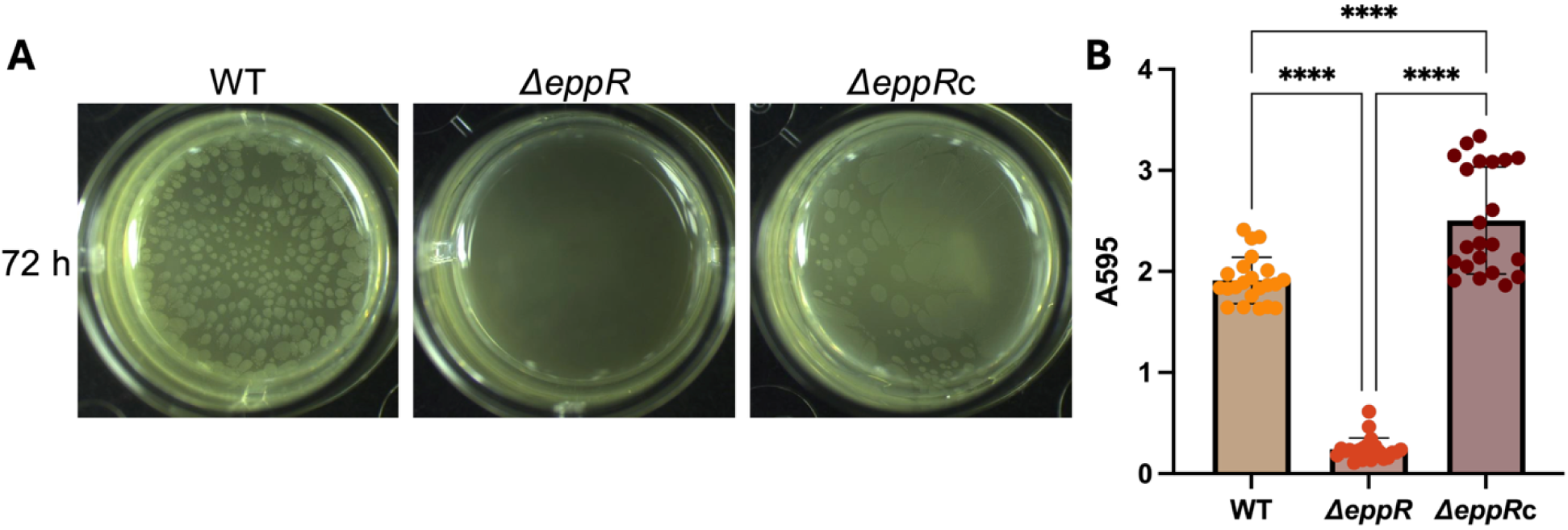
EppR is required for cell surface attachment and biofilm formation. (A) Knockout of *eppR* results in loss of biofilm formation. Pellicle biofilms of WT, Δ*eppR*, and Δ*eppR* complemented with *eppR* under its own promoter on a pNLAC plasmid (Δ*eppR*c) after 72 h of growth. WT and Δ*eppR* strains contain empty pNLAC plasmid. Representative images are shown. (B) Cell attachment to an abiotic surface is significantly reduced in Δ*eppR.* Cell attachment to polystyrene was quantified by Crystal Violet staining by measuring absorbance at A595. At least 20 replicates were measured for each strain over three independent experiments. Statistical analysis was performed using one-way ANOVA with multiple comparison by Tukey-test. **** indicates a p-value of < 0.0001.

### EppR is involved in the regulation of some biofilm-related genes

Up to this point we have shown that *eppR* is somehow involved with biofilm formation. To learn of a possible mechanism, we decided to use a genome-wide approach and followed these experiments with RNAseq to assess the transcriptome of the Δ*eppR* strain compared to WT (Figure 2). We chose to perform the RNAseq in exponentially grown cells to learn about all the possible changes caused by lack of EppR. We surmised that at minimum we should detect differential expression in biofilm genes, identified by our group (Ching et al., 2019, 2024) or elsewhere (Ronish et al., 2019; Tomaras et al., 2003, 2008) in the Δ*eppR* compared to the WT strain.

**Figure 2:**
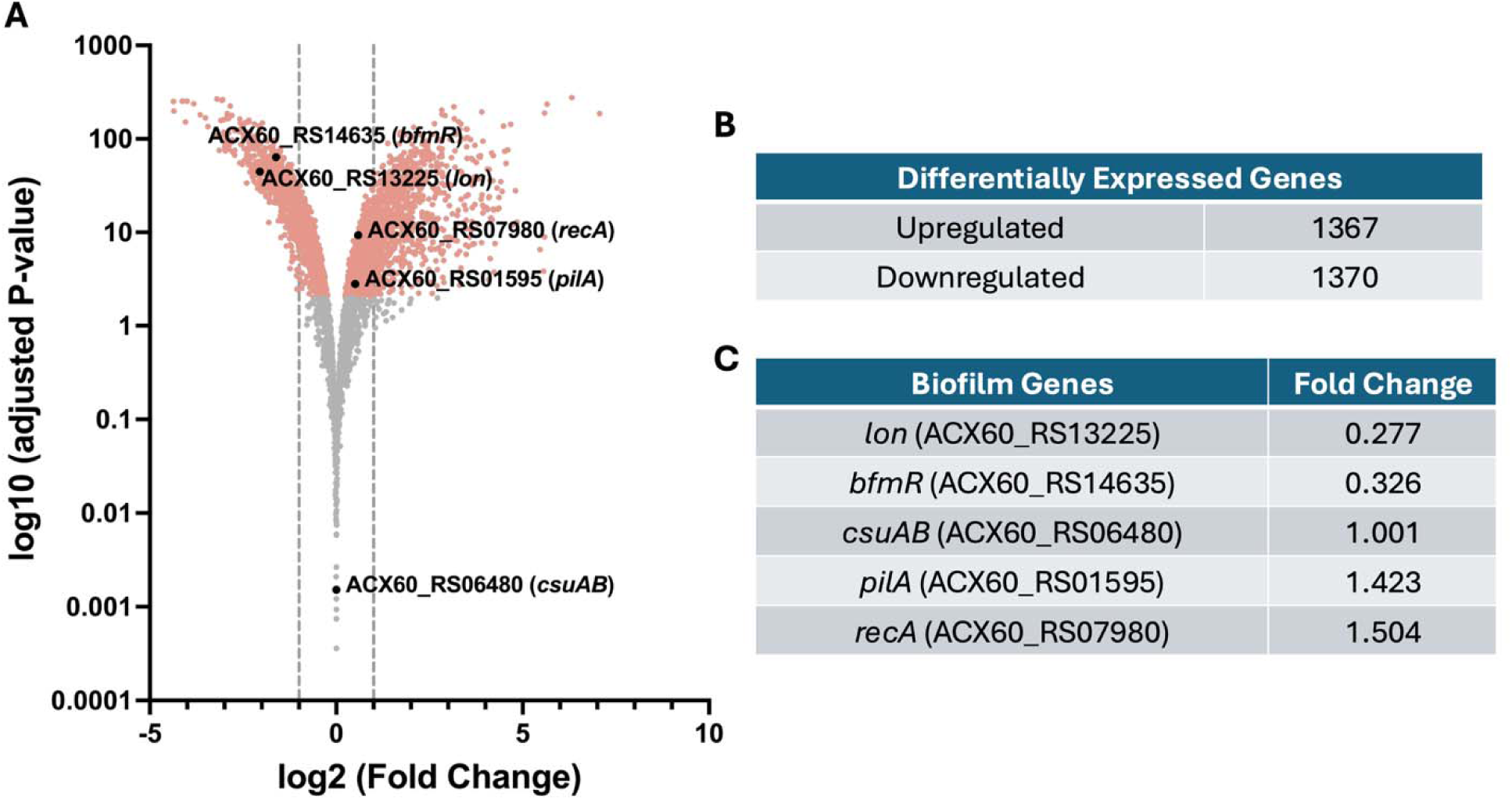
Global transcriptome analysis of Δ*eppR* compared to WT show large number of differentially expressed genes including key biofilm regulators. (A) Volcano Plot depicts differentially expressed genes based on RNAseq data from exponential phase cells. Red dots denote genes that have a log (adjusted P-value) > 2, which indicates changes in gene expression are significant. Fold change (FC) filters of - 1<Log2(FC)<1 was omitted in this study to identify differences in genes that typically have low expression, though are depicted by the gray dotted lines. Genes of interest in this study are denoted by the black dots. (B) Table showing the total number of upregulated and downregulated genes in Δ*eppR* compared to WT. (C) Table of key biofilm genes investigated in this study with their respective changes in expression.

Genes were considered differentially expressed if they had an adjusted p-value of <0.05, which would indicate that the changes in gene expression are significant. Fold change filters were not used to identify differences in genes that typically have low expression (Cook et al., 2023).

Initial analysis of RNAseq data shows a surprisingly large number of differentially expressed genes in Δ*eppR,* affecting the expression of a wide range of genes, a pattern typical of a global transcriptional regulator (Deyell et al., 2025; Shen-Orr et al., 2002). We identified ∼1300 upregulated genes and about the same number of downregulated genes (Figure 2, Table S1). Even when a fold change (FC) filter is applied for values 0.5 < FC < 2, ∼800 genes are upregulated and ∼700 genes are downregulated, indicating that a global transcriptional regulator was affected.

For this study, we focused on genes previously shown to be involved in biofilms, such as *recA* (A1S_1962, ACX60_RS07980), *bfmR* (A1S_0748, ACX60_RS14635), *csuAB* (A1S_2218, ACX60_RS06480), *pilA* (A1S_3177, ACX60_RS01595), and *lon* (A1S_1030, ACX60_RS13225) (Ching et al., 2019, 2024; Harding et al., 2013; Ronish et al., 2019; Tomaras et al., 2003, 2008; Vesel & Blokesch, 2021). RecA, the cells’ main recombinase, regulates the DDR and is involved in biofilm regulation (Ching et al., 2024; Norton et al., 2013). BfmR was defined as the central biofilm regulator (Tomaras et al., 2008). *csuAB* encodes subunits of the Csu pili, which mediates cell attachment primarily on abiotic surfaces (Ahmad et al., 2023; Tomaras et al., 2003), and *pilA* encodes the major subunit of type IV pili, involved in twitching motility and natural competence, as well as adherence to epithelial cells (Harding et al., 2013; Ronish et al., 2019; Vesel & Blokesch, 2021). Lon is a protease somehow involved in biofilm formation likely by regulating changes in the cell envelope (Ching et al., 2019).

We find that in Δ*eppR* there is both modest increase in *recA* expression and a large decrease in *bfmR* expression. This is consistent with previous findings in our lab in which cells containing higher levels of *recA* make less biofilm and have lower expression of *bfmR* (Ching et al., 2024). We see that though there is a modest increase in *pilA* expression (Figure 2C), this may not be remarkable as we know that *pilA* is not heavily involved in the attachment to abiotic surfaces (Ahmad et al., 2023). However, our RNAseq data show that there is no significant change in *csuAB* expression, which is inconsistent with the data in Figure 1. We find that there is a significant decrease in *lon* expression in Δ*eppR*, consistent with previous findings showing that strains lacking *lon* are deficient in biofilm formation (Ching et al., 2019). While the expression of these genes in the RNAseq data are mostly consistent with the idea that *eppR* is needed for biofilm formation, the expression of the adhesins PilA and Csu encoding genes seems to conflict with this hypothesis. Because we chose to perform the RNAseq on exponentially grown cells, we figured that *pilA* and *csu* expression maybe mostly consistent with exponential growth and not biofilms. Therefore, we performed additional experiments in biofilm cells.

### EppR affects pili gene expression

We decided to measure the expression of the adhesin genes, *pilA* and *csuAB*, in biofilm conditions. We grew WT, Δ*eppR*, and Δ*eppR*c biofilms for 72 h in 24-well plates, collected all cells in the well, and extracted RNA to perform qPCR. We detected a ∼4- fold decrease in *pilA* expression in Δ*eppR* and an ∼2-fold increase in Δ*eppR*c (Figure 3A) suggesting that PilA may play a role in adhesion, though unclear (Chabane et al., 2011; Eijkelkamp et al., 2011), and agreeing with the idea that EppR has a role in biofilm formation.

**Figure 3:**
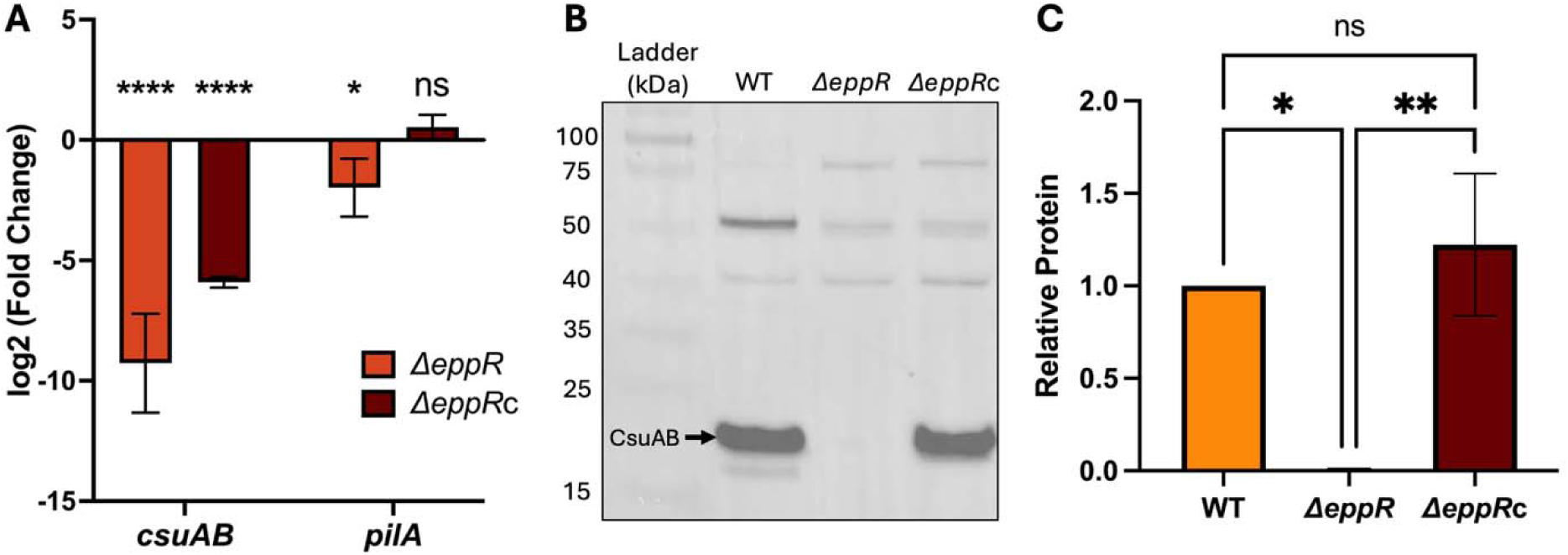
EppR affects the production of key surface adhesion pili. (A) Quantitative RT-PCR experiments on biofilm cells show downregulation of key pili genes, *csuAB* and *pilA*, in Δ*eppR.* Y-axis denotes Log2 fold change relative to WT. qRT-PCR experiments were performed in biological and technical triplicate. Ordinary two-way ANOVA with multiple comparison by Tukey-test was used for statistical analysis. Statistical markers are derived from pairwise comparison to WT. * indicates a p-value of < 0.05, **** indicates a p-value of < 0.0001, and ns indicates not significance. (B) Western blot for CsuAB on biofilm cells shows Δ*eppR* cells do not make Csu pili. Δ*eppR*c shows similar levels of CsuAB compared to WT. CsuAB is denoted by the arrow. Image is representative of four independent experiments. Antibodies used and their respective dilutions are indicated in Materials and Methods. (C) Quantification of the blot shown in (B) shows that Δ*eppR* produces little to no CsuAB. Quantification was performed by Fiji. Ordinary one-way ANOVA with multiple comparison by Tukey-test was used for statistical analysis. * Indicates a p-value of < 0.05, ** indicates a p-value of < 0.01, and ns indicates not significant.

As expected, we found that in Δ*eppR*, there was an ∼500-fold decrease in *csuAB* expression when compared to WT, consistent with the phenotypic biofilm data (Figure 1), especially since Csu pili are the main pili used to adhere to abiotic surfaces (Ahmad et al., 2023; Tomaras et al., 2003). Interestingly, while Δ*eppR*c makes similar biofilms compared to WT (Figure 1), we detected an ∼60-fold decrease in *csuAB* expression compared to WT though still exponentially higher expression than Δ*eppR*. While the expression of *csuAB* in Δ*eppR*c is not at the WT levels, it may be sufficient to produce enough adhesin for biofilm attachment. These data provide evidence that the lack of biofilm seen in Δ*eppR* could be attributed to reduced expression of the adhesin-coding genes.

Since we found that Δ*eppR*c makes similar biofilms to WT despite having lower *csuAB* expression, we next investigated CsuAB protein levels in the biofilms. To do this, we grew WT, Δ*eppR*, and Δ*eppR*c biofilms, collected all cells in the wells, extracted proteins, normalized protein concentrations and performed a western blot using an antibody specific for CsuAB (Figure 3B and C) (Tomaras et al., 2008). We found that, despite the lower expression found by qPCR, Δ*eppR*c and WT had similar levels of CsuAB protein at the same total protein concentrations tested. This suggests that at comparable cell concentrations the levels of *csuAB* expression in Δ*eppR*c are sufficient to produce enough CsuAB adhesins to form biofilms. In Δ*eppR*, as expected and consistent with the qPCR data, CsuAB was decreased, almost below the limit of detection. The data provide additional evidence to the finding that Δ*eppR* lack of biofilms is due to reduced pili. We then surmise that both qPCR and western blot data from biofilms support the idea that EppR is involved in the regulation of biofilm formation as we observe lower pilus production when *eppR* is deleted.

### EppR affects RecA expression

Since previous work in our lab demonstrated that RecA levels modulate Csu pili (Ching et al., 2024) and we have shown here that the deletion of EppR results in lower levels of Csu pili, we next investigated a link between EppR and RecA. We also noticed that in the RNAseq data (Figure 2), there is a slight, but significant increase of *recA* expression in Δ*eppR* compared to WT consistent with the known anti-correlation of high RecA and low biofilms (Ching et al., 2024).

We next performed an additional qPCR experiment measuring *recA* expression in the WT compared to Δ*eppR* (Figure 4A). The obtained qPCR data is largely consistent with the RNAseq; an increase in *recA* expression in Δ*eppR* compared to WT, though we measured a greater increase in expression in the qPCR (∼4-fold) compared to the RNAseq data (∼1.5-fold). Additionally in the qPCR, Δ*eppR*c *recA* expression was like WT. This data provides evidence that EppR somehow influences *recA* expression.

**Figure 4:**
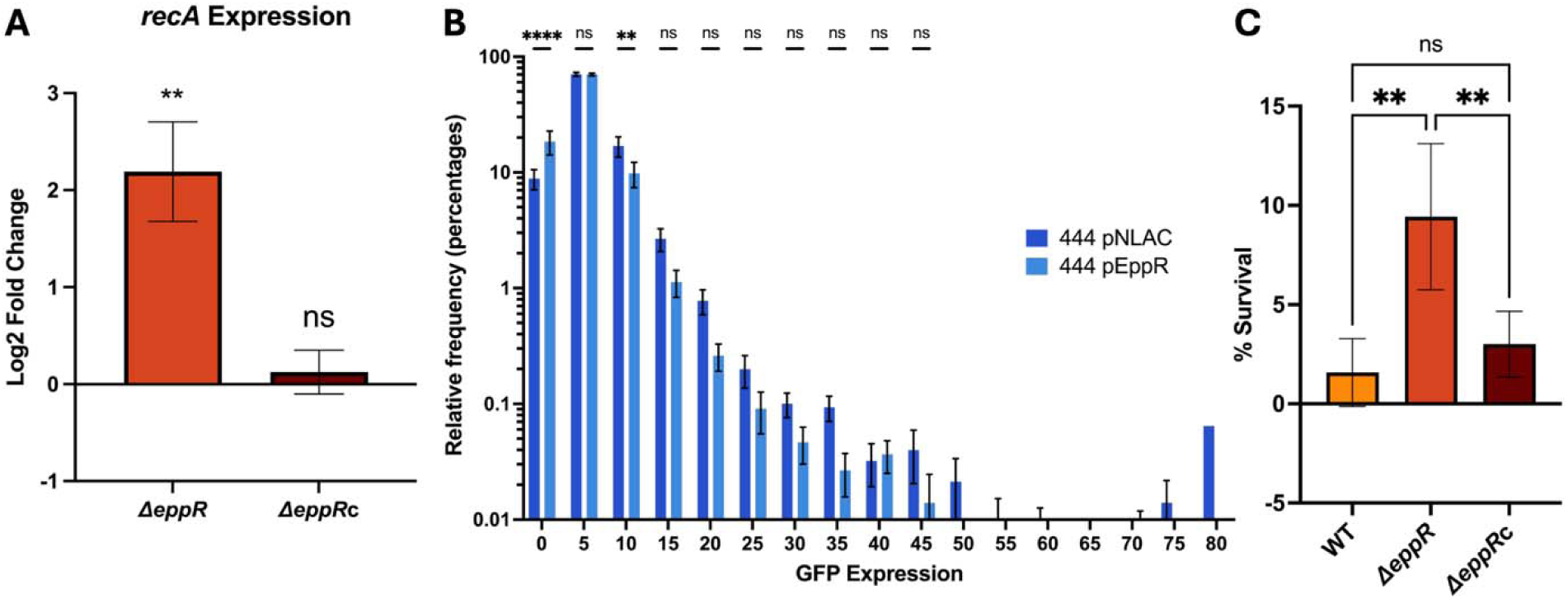
EppR influences *recA* expression. (A) Quantitative RT-PCR experiment shows deletion of *eppR* results in a significant upregulation of *recA* expression compared to WT. Log2 fold change relative to WT. qRT-PCR experiments were performed in biological and technical triplicate. Ordinary two-way ANOVA with multiple comparison by Tukey-test was used for statistical analysis. Statistical markers are derived from pairwise comparison to WT. ** indicates a p-value of < 0.01 and ns indicates not significance. (B) Overproduction of EppR results in a significant increase in a subpopulation of cells with little to no *recA* expression. Single cell fluorescence microscopy shows a greater percentage of cells that have very low or no *recA* expression measured as fluorescence from a RecA-GFP fusion protein on the chromosome (444). The fluorescence of 444 with empty pNLAC is compared to the strain with pEppR, (pNLAC + *eppR* under its own promoter). The GFP expression was binned and the percentage of cells within each bin was graphed. The Y-axis is shown on log scale to better visualize the differences between the two strains. At least 10,000 cells from 9 independent experiments were quantified using Fiji (Schindelin et al., 2012). Ordinary one-way ANOVA with multiple comparison by Tukey-test was used for statistical analysis. ** indicates a p-value of < 0.01, **** indicates a p-value of < 0.0001, and ns indicates not significant. (C) Deletion of *eppR* results in increased resistance to UV. Cells were exposed to 150 J/m^2 UV light. Three independent experiments were performed. Ordinary one-way ANOVA with multiple comparison by Tukey-test was used for statistical analysis. ** indicates a p-value of < 0.01 and ns indicates not significant.

To look further into the effect of EppR on RecA, we surmised that elevating EppR levels in cells would result in low cellular RecA. To test this hypothesis, we introduced pEppR (pNLAC plasmid (Luke et al., 2010) containing the *eppR* gene under its own promoter; Table 1) into a strain that contains a *recA* – *gfp* translational fusion in the *A. baumannii* genome (444; Table 1) (Ching et al., 2017). This strain has a native *recA* gene in addition to the *recA – gfp* fusion. We then performed single cell fluorescence microscopy of cells with and without extra copies of EppR and quantified *recA* expression based on the GFP fluorescence using Fiji (Schindelin et al., 2012). We counted at least 10,000 single cells with and without the pEppR plasmid. The average fluorescence from the 444 cells with the empty pNLAC plasmid is like that of the 444 with pEppR. This data would indicate that increasing EppR doesn’t necessarily downregulate *recA* expression (Figure S2). However, while the scatter plot shows no large differences in average RecA levels, we detect a larger subpopulation of cells with lower GFP expression in 444 with pEppR that is absent in 444 pNLAC. To better visualize this difference, we graphed the frequency of cells with low and high fluorescence by binning RecA-GFP fluorescence (Figure 4B). We find a significantly higher frequency of cells in 444 pEppR with little to no *recA* expression (GFP expression ≈ 0) compared to the control. In addition, cells with the empty pNLAC plasmid display higher frequency of fluorescent cells with values ≥ 10. This suggests that while high EppR doesn’t seem to reduce, on average, the levels of cellular RecA, a higher frequency of cells with low RecA is detectable only in cells with additional copies of EppR. We have seen this before in previous work in our lab, where in response to DNA damage, heterogeneity in RecA and other genes is apparent (MacGuire et al., 2014).

Lastly, to provide additional evidence that EppR influences RecA levels, we performed an ultraviolet (UV) sensitivity assay (Figure 4C), as it is known that low RecA results in increased sensitivity to UV, as RecA is necessary for *A. baumannii* to survive DNA damage (Aranda et al., 2011). We hypothesized that due to increased RecA in Δ*eppR*, we should observe comparatively greater resistance to UV. When exposed to 150 J/m^2^ of UV light, a level recommended for screening mutations in RecA (see Materials and Methods), Δ*eppR* cells have greater resistance to UV than WT (Figure 4C), as there was a 10-fold increase in the percentage of cells that survived UV treatment. When *eppR* is complemented in Δ*eppR*c, we see that the percentage of survivors is reduced to levels like WT. With these experiments, we have provided evidence that EppR somehow regulates *recA* through unknown mechanisms.

To investigate possible mechanisms of regulation, we tested whether EppR could bind to the *recA* regulatory region, which includes the *recA* promoter and a 100 bp DNA sequence that encodes a 5’ untranslated region (UTR), that we have previously shown to influence intracellular RecA protein levels (Ching et al., 2017). We have also shown that EppR represses genes that encode DNA polymerase V through the direct binding of a conserved motif present in their promoter sequences (Nguyen et al., 2025).

We then amplified the *recA* regulatory region by PCR using fluorescent primers (Table 2). The labeled fragment was used in electrophoretic mobility shift assays (EMSA) by incubating purified EppR (Nguyen et al., 2025) with the fluorescent-labeled probe. Products were separated in a non-denaturing polyacrylamide gel (Figure S3). We found that EppR binds to this regulatory region, with full binding observed at ∼1600 nM of EppR (Figure S3A). Binding data was quantified using Fiji (Schindelin et al., 2012), and the dissociation constant (K_D_) of this binding reaction was calculated at ∼1100 nM (Figure S3B). The ability of EppR to bind to the *recA* regulatory region is much weaker than its ability to bind to its own promoter, where the K_D_ is ∼10-fold lower (K_D_ ≈120 nM) (Nguyen et al., 2025). After observing that EppR binds to the *recA* regulatory region, we searched for a potential binding motif. We analyzed the *recA* regulatory region with the Motif for Em Elicitation (MEME) program (Bailey et al., 2009), using the EppR binding motif (Nguyen et al., 2025). We identified an inverted repeat within the 5’-UTR sequence that could potentially serve as the EppR binding site (Figure S3C). While this sequence in the *recA* 5’-UTR was detected by MEME as the best match, there are significant differences to the conserved EppR binding motif, which is consistent with the relatively weak binding affinity shown in the EMSA. With this data, we provide evidence that EppR may regulate *recA* through direct repression, though the relatively weak binding shown in the EMSA suggest there may be additional RecA regulators that have yet to be identified.

**Table 2:**
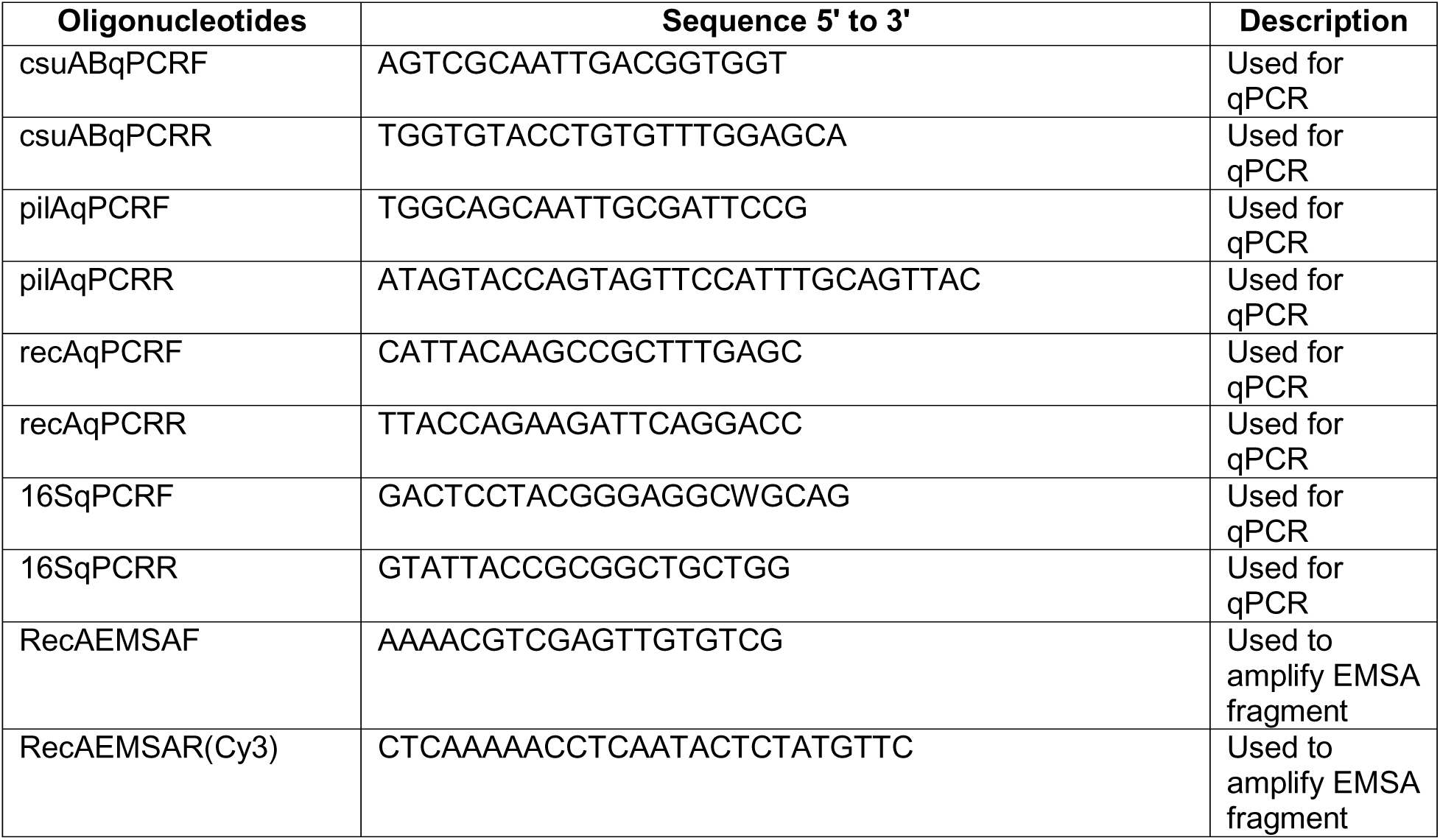
Oligonucleotide List.

### EppR interacts with RecA during biofilm conditions

Lastly, since we have shown that EppR influences RecA levels perhaps by affecting *recA* expression, we wanted to look at RecA protein levels under biofilm conditions. Using an antibody specific for RecA that we have previously worked with (Ching et al., 2017, 2024; Norton et al., 2013), we performed a western blot on proteins collected from cells in biofilms (Figure 5A), which separated into soluble and insoluble fractions. In WT, we detected two bands, one of ∼40 kDa, which corresponds to the RecA monomer (∼39 kDa) in the insoluble fraction and another one between the 50 and 75 kDa size markers in the soluble fraction. Previous work using this antibody (Ching et al., 2024; Norton et al., 2013) had only detected the RecA monomer band in exponential phase cells, indicating that the larger band may be only present in biofilm conditions. It is possible that the protein in the larger band is RecA in complex with another protein. Though SDS-PAGE is expected to separate complexes, it is not rare to see them (Cafarelli et al., 2013). It is unlikely that this band corresponds to a RecA dimer as its size would be ∼80 kDa. In Δ*eppR*, there is a RecA band in the insoluble fraction and no band in the soluble fraction corresponding to the putative complex, which is complemented in Δ*eppR*c, where the band intensity is like WT. A potential complex made up of RecA and EppR (∼24 kDa) would have a size of ∼63 kDa, which would be consistent with the size of this larger band, providing preliminary evidence that EppR complexes with RecA in biofilms.

**Figure 5:**
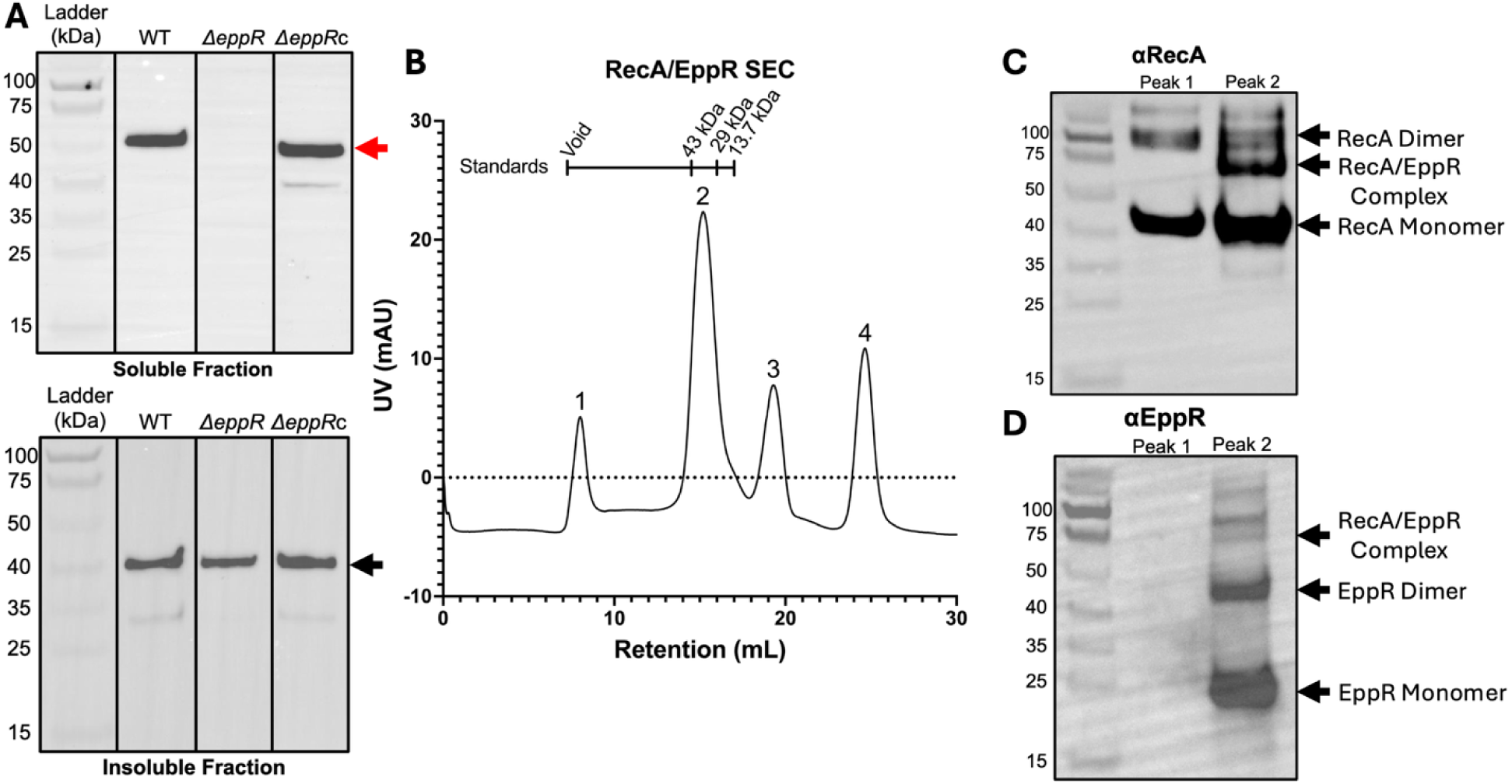
EppR interacts with RecA in biofilms. (A) RecA detection in biofilm cells. Western blots with αRecA antibody were used to detect RecA from both soluble and insoluble protein fractions. A band between 50 and 75 kDa, consistent with a RecA- EppR complex, shown by the red arrow in the soluble fraction (top image) is present in WT and Δ*eppR*c (biofilm proficient) and absent in Δ*eppR* (biofilm deficient). Monomeric RecA levels are similar in all strains as shown by the black arrow in the insoluble fraction (bottom image). Images are representative of three independent experiments. (B) Size exclusion chromatography (SEC) of RecA mixed with EppR results in four distinct peaks. (C) Western blot probing for RecA shows RecA monomers and multimers in Peak 1. Peak 2 shows both RecA monomer and multimers along with the potential RecA/EppR complex. (D) Western blot probing for EppR shows no EppR in Peak 1 while Peak 2 contains an EppR dimer and monomer, along with a band consistent with a RecA/EppR complex. At the settings used to develop the blots, neither EppR nor RecA were detected in Peaks 3 and 4 as the concentration of protein within those peaks were below the limit of detection. Antibodies used and their respective dilutions are indicated in the Materials and Methods.

To provide evidence that this complex exists, we mixed purified EppR and purified RecA and performed size exclusion chromatography (SEC) (Figure 5B). We hypothesized that there should be a fraction where both proteins elute together. The resulting SEC separation of the EppR-RecA mixture displayed four defined peaks (Figure 5B). To determine the identity of the proteins in each peak, we separated these respectively by SDS-PAGE and performed western blots using αRecA and αEppR antibodies (Figure 5C and D, respectively). We observed that the first peak consisted of only RecA; EppR was not detected. In the second peak, we see that both RecA and EppR are present. This data alone does not provide foolproof evidence of the presence of the complex as we have shown before that the EppR dimer (with a mass of ∼48 kDa) elutes at a similar time frame in the same column (Nguyen et al., 2025). However, in addition to only RecA and EppR, we detected a strong band in the αRecA blot that is separated at a mass like an EppR/RecA complex (∼63 kDa; Figure 5C). In the αEppR blot, we also observe a band around the same mass, providing evidence that a complex between the two proteins is likely. We did not detect either protein in peaks 3 or 4; perhaps these are at very low concentrations. These results indicate that in biofilm cells RecA complexes with EppR and this complex is somehow needed for biofilm formation. A possible complex between EppR and RecA was modeled by Alphafold 3 (Abramson et al., 2024). In this putative complex, EppR would bind to the amino end of RecA (Figure S5). More research is currently being conducted in the lab to learn about this potential complex.

## Discussion

Previous work has shown that in many bacterial species, there is a DDR-biofilm axis (Costa et al., 2014; Gozzi et al., 2017; Inagaki et al., 2009; Linares et al., 2006; Takahashi et al., 1995; Walter et al., 2015). The same hold true for *A. baumannii*. The levels of RecA, a key DDR regulator, influences biofilm development, where low RecA results in high biofilm and high RecA results in little to no biofilm formation (Ching et al., 2024). In this work, we expand on the mechanistic understanding between the DDR and biofilm development.

We recently identified a novel TetR-family transcriptional regulator, EppR, and shown it to be a direct repressor of genes encoding the regulator UmuDAb and of DNA polymerase V (DNA Pol V) genes encoding error-prone DNA polymerases used in translesion synthesis (Reuven et al., 1999; Tang et al., 1999; Wang, 2001). EppR and UmuDAb regulate each other and together they contribute to higher expression of DNA Pol Vs which in turn contribute to the rise of antibiotic resistance through the mutagenesis of drug targets (Boshoff et al., 2003; Cirz et al., 2005; Cirz, O’neill, et al., 2006; Cirz & Romesberg, 2006). Here, we find that EppR is also involved in the regulation of biofilms. We find that an *eppR*-deficient strain does not form biofilms (Figure 1). RNAseq data comparing global gene expression in Δ*eppR* to WT show decreased *bfmR* expression as well as an increase in *recA* (Figure 2). These data are consistent with previous studies in our lab and others that had identified these genes as important for biofilm development (Ching et al., 2019, 2024; Tomaras et al., 2008). We also show that lack of biofilms in Δ*eppR* is likely due to downregulation of adhesin genes (Figure 3), namely *csu*, encoding the archaic Csu pili, that mediates cell attachment to abiotic surfaces. Finally, we provide a potential mechanism by which EppR modulates biofilms; EppR influences *recA* expression, and loss of EppR results in increased *recA* expression (Figure 4) and that may be due to EppR acting as a direct *recA* repressor (Figure S3). Moreover, we provide evidence that, in biofilm cells, EppR is in complex with RecA, to further control RecA levels.

TetR-family transcriptional regulators are typically known as regulators of antibiotic efflux pumps, though they have been found to be involved in a variety of cellular processes, such as cell-cell signaling, cell division, and metabolism (Cuthbertson & Nodwell, 2013; Ramos et al., 2005). Despite being involved in many different roles, EppR is the first TetR-family regulator involved in the DNA damage response (Nguyen et al., 2025). Regulation of biofilms by TetR-family proteins has been defined in other bacterial species such as *Staphylococcus, Streptococcus* and *Vibrio* species, though their roles vary (Cho et al., 2002; Conlon et al., 2002; Cramton et al., 1999; Croxatto et al., 2002; Liu et al., 2017; McDougald et al., 2001). Like our data in Figure 1, it has been shown in *Vibrio anguillarum* that a homologue of *Vibrio harveyi* LuxR, VanT, is needed for biofilm formation, as a *vanT*-deficient strain is unable to form biofilms (Croxatto et al., 2002). Interestingly, another *V. harveyi* LuxR homologue in *Vibrio vulnificus*, SmcR, had the opposite effect. A *smcR* mutant strain has increased biofilm formation compared to a *smcR^+^* strain (McDougald et al., 2001). Like SmcR, a deletion of the TetR-family regulator BrpT in *Streptococcus sanguinis* results in increased biofilm formation, due to increased filamentous structures in the biofilm and increased extracellular glucans (Liu et al., 2017). In both *Staphylococcus aureus* and *S. epidermidis*, *icaR* encodes a TetR-family regulator of the *ica* locus, which in turn encode intracellular adhesins (Cho et al., 2002; Conlon et al., 2002; Cramton et al., 1999). It has been shown in both species that the *ica* locus is required for biofilm formation (Cho et al., 2002; Cramton et al., 1999), though the role of *icaR* is not as straightforward. In the presence of ethanol, *icaR* is downregulated, leading to the activation of the *ica* locus and biofilm formation (Conlon et al., 2002). Interestingly, the same study investigated *icaR* transcription in a high salt environment, which can promote biofilms in *S. epidermidis* (Rachid et al., 2000), but saw no change in expression. This study, along with others, has shown not only that TetR-family transcriptional regulators are involved in biofilm development, but highlights how diverse their roles can be.

In addition to modulating the levels of adhesins in cells, EppR appears to regulate *recA* expression. We have previously shown that low RecA levels result in high biofilm formation (Ching et al., 2024). We find that EppR can bind to the *recA* regulatory region, which includes the *recA* promoter and a 5’-UTR, which we have shown influences *recA* expression (Figure S2) (Ching et al., 2017). Both RNAseq and qPCR data agree that the loss of *eppR* results in an upregulation of RecA, leading to reduced biofilm formation (Figure 2 and 4). It seems likely that through regulating RecA levels, EppR influences *csu* expression, as we have previously characterized a pathway where low RecA results in high levels of another DDR regulator UmuDAb (A1S_1389; ACX_RS11175) that functions as an activator of the *bfmR* gene, encoding BfmR needed for induction of the *csu* operon and thus biofilm formation (Ching et al., 2024; Tomaras et al., 2008).

In addition to transcriptional data, we find that biofilm cells of Δ*eppR* lack a strong band recognized by an αRecA antibody located between the 50 and 75 kDa size markers that is present in WT and the Δ*eppR* complement strain (Figure 5A). Due to the high specificity of the RecA antibody (Ching et al., 2017; Norton et al., 2013), it is likely that this band consists of RecA in complex with another protein despite these being separated in SDS-PAGE, as we have seen this before (Cafarelli et al., 2013). Based on the size of the band and the fact that it is not present in Δ*eppR*, it seemed likely that this is a complex between a monomer of RecA and a monomer of EppR, which would result in ∼63 kDa. When we investigated this further by performing SEC on a mix of purified EppR and RecA, we detected a band around the expected mass with both EppR and RecA antibodies (Figure 5B and C). Interestingly, levels of monomeric RecA are unchanged between Δ*eppR* and WT, indicating that this complex may be needed to promote biofilm formation, though more studies would need to be done to confirm this. These data show that EppR can influence biofilms by affecting RecA, not only at the transcriptional level, but also post-translation.

Lastly, this study provides evidence that EppR is involved in cellular processes beyond biofilm development and the DDR. In the RNAseq data, we see that the deletion of *eppR* results in the downregulation of both *lon* and *bfmR* (Figure 2). While we and others have shown that both proteins are needed for biofilm formation (Ching et al., 2019; Tomaras et al., 2008), these proteins have functions that are not just limited to biofilms. *lon* encoding Lon protease, is primarily involved in targeted degradation of misfolded proteins and certain regulators (Charette et al., 1981; Fu et al., 1997; Goldburg, 1992). In *A. baumannii*, we observed that Lon is involved with UV resistance, motility, and capsule production (Ching et al., 2019). While BfmR was initially identified as a biofilm regulator, a recent study showed that it is implicated in a variety of cellular processes (Raustad et al., 2025). It was shown that BfmR has a role in virulence, acquisition of antibiotic resistances and its regulon includes genes involved in capsule production, protein folding, production of outer membrane proteins, and type IV pilus production. We show that EppR can also influence the expression of type IV pilus genes (Figure 3). While type IV pili have been shown to promote adherence to biotic surfaces (Piepenbrink et al., 2016), type IV pili are also required for twitching motility and natural competence (Harding et al., 2013). Remarkably, the RNAseq data show that the knockout of *eppR* results in the differential expression of over 2600 genes, indicating that the function of EppR is widespread in cells.

In summary, these findings further characterize the regulation of biofilm development in *A. baumannii* and solidify the link between biofilm development and the DNA damage response.

### Strains and Culture Conditions

All bacterial strains (Table 1) were grown in Miller-LB medium (10 g L^−1^ Tryptone, 5 g L^−1^ Yeast Extract, 10 g L^−1^ NaCl) at 37°C, with shaking at 225 rpm unless indicated otherwise. Solid medium contained 1.5% agar (Fisher Bio-Reagents). Antibiotics were added to the medium at the indicated concentrations unless stated otherwise. Kanamycin (Km, 35 μg/mL), carbenicillin (Carb, 100 μg/mL), tetracycline (Tet, 6 μg/mL), and gentamicin (Gm; 10 μg/mL).

### Electroporation of Plasmids into *A. baumannii*

Plasmids were introduced into *A. baumannii* as previously described (MacGuire et al., 2014). Briefly, competent cells were prepared by growing strains to exponential phase in Miller-LB broth. Cells were collected by centrifugation and washed multiple times in sterile deionized water. Cells were then resuspended in 20% glycerol. For transformations, 50 μL of competent cells were mixed with 50 ng of the plasmid of interest and transferred into an aluminum-coated electroporation cuvette with a 1 mm gap. A Bio-Rad Micropulser with standard settings for *Escherichia coli* (1.8 kV) was used for electroporation.

### Quantification of Biofilm with Crystal Violet

Quantification of biofilms were performed as previously described (Brychcy et al., 2024). Briefly, pellicle biofilms are set up in multi well dishes, incubated, washed and stained with Crystal Violet which is then dissolved in 30% acetic acid and read in a spectrophotometer. The intensity of the stain directly correlates with biofilm strength. Strains were inoculated 1:1000 in YT media (10 g L^−1^ Tryptone, 1 g L^−1^ yeast extract, 2.5 g L^−1^ NaCl). Cultures were grown without agitation for 72 h at 25°C on a non-tissue-culture treated 24 well plate. Cultures were then collected through aspiration and wells washed with PBS and dried in a biosafety cabinet for 10 min. The dried wells were stained with 0.1% Crystal Violet for 15 min at room temperature. The Crystal Violet solution is discarded, and wells are washed twice with PBS. 30% glacial acetic acid was used to dissolve the remaining dye. The plate was incubated at room temperature while shaking at 120 rpm for 12 min before measuring A595 in an Agilent BioTek Epoch 2 plate reader. Measurements were performed at least three independent times. Biofilm images were taken with a Leica MSV296 dissecting microscope and a Leica DMC2900 camera.

### RNAseq Experiment and Analysis

The *A. baumannii* ATCC 17978 WT and isogenic Δ*eppR* strains were grown to exponential phase in Miller-LB medium and ∼10^8^ cells were collected and sent to Genewiz for RNA extraction, rRNA depletion, RNA fragmentation, quality control, library preparation, and sequencing through Illumina HiSeq 2x150bp. Three biological isolates from each strain were sent for RNAseq. Raw sequencing reads were uploaded to the National Center for Biotechnology Information (NCBI, BioProject: PRJNA1272101). Sequence quality was assessed using FastQC and reads were trimmed with Trimmomatic. Reads were aligned to the ATCC 17978 genome (NZ_CP012004) using STAR and differential gene expression was analyzed with DESeq2 (Bioconductor). Genes with an adjusted P value < 0.01 were considered differentially expressed. To identify differences in genes that typically have low expression, fold change filters were not applied. Volcano plots were created using Graphpad Prism.

### Quantitative PCR

Strains were diluted 1:1000 from a saturated culture in fresh Miller LB broth and grown for three hours until mid-exponential phase was reached. Total RNA was extracted, and cDNA was synthesized as previously described (Brychcy et al., 2024). qPCR was performed with Luna® Universal qPCR mix (New England Biolabs) in QuantStudio™ 3 (Applied Biosystems). All procedures were performed according to the manufacturer’s instructions. The ΔΔCt method with normalization against 16S RNA was used to determine relative gene expression. Primers were designed using PrimerQuest™ (Integrated DNA Technologies) and optimized for a melting temperature of 60°C. Primers used are listed in Table 2. Each measurement was performed in both technical and biological triplicate.

### Detection of RecA, CsuAB, and EppR in Biofilms

RecA, CsuAB, and EppR proteins were detected by Western blot. Strains were diluted from a saturated culture 1:1000 in YT media (10 g L^−1^ Tryptone, 1 g L^−1^ yeast extract, 2.5 g L^−1^ NaCl). Cultures were grown without agitation for 72 h at 25°C on a non-tissue-culture treated 24 well plate. All cells within the wells were collected and whole protein lysates were prepared using Bugbuster reagent (Novagen) and Pierce Universal Nuclease (Thermo Fisher) according to the manufacturer’s instructions. Cell-free lysates were quantified by Bradford assay following the manufacturer’s instructions. Samples were normalized by protein mass and mixed with Laemmli buffer. Samples were then incubated at 100°C for 10 min, before separation in a 12% polyacrylamide SDS gel in MES buffer (Invitrogen). Protein samples were transferred to a methanol-activated PVDF membrane as previously described (Norton et al., 2013). Membranes were exposed to the following primary antibodies: anti-RecA (1:10,000) (Abcam), anti- CsuA/B (1:10,000) (Tomaras et al., 2008), and anti-EppR (1:5,000) (custom made by Abclonal), and followed by a 1:10,000 dilution of an Anti-rabbit-HRP secondary antibody (Abcam). Bands were visualized using chemiluminescence via SuperSignal West Pico Plus (Thermo Fisher) and imaged with a BioRad Chemidoc MP. Samples were run in parallel in a second gel and stained with Coomassie Brillant Blue to confirm equal loading of protein. Ladder used is the Spectra^TM^ Multicolor Broad Range Protein Ladder (Thermo Scientific). Fiji (Schindelin et al., 2012) was used to quantify relative protein levels.

### Fluorescence Microscopy

Strains were diluted 1:1000 in fresh Miller-LB broth from saturated cultures and grown until mid-exponential phase was reached. 2 µL of each were placed on a 1% agarose in water pad and covered with a coverslip before imaging. Single cells were visualized using a Leica MicroStation 5000 with a Leica DM3000G camera (Leica) at 100X magnification. All images were taken in phase contrast or with a GFP filter (Excitation: 450 nm – 490 nm, Dichromatic mirror: 495 nm; Emission: 500 nm – 550 nm). Exposure was set to 200 ms. Mean fluorescence was quantified using Fiji (Schindelin et al., 2012) with a custom script that was previously used (Brychcy et al., 2023). The custom script opens both phase contrast and fluorescent images and converts them to 16-bit images. To identify cells, the phase contrast image is converted to black and white using the ‘Threshold’ function of Fiji. Connected cells were separated using the ‘Watershed’ function. Fluorescence intensity was measured using the ‘Analyze Particles’ function, referencing the fluorescent image for intensity. All data was compiled, analyzed, and displayed in graphs using Graphpad Prism.

### UV Survival Assay

UV survival assays were performed as previously described (Ching et al., 2017). Saturated cultures were diluted 1:100,000 in PBS and 50 μL of this dilution was evenly spread using sterile glass beads on LB agar plates. Plates were irradiated using a Stratalinker 1800 UV Crosslinker (Stratagene) with 150 J/m^2^, as it was the manufacturer instruction for screening of Δ*recA* bacterial strains. Colonies on untreated and UV-treated plates were counted, and percent survival was standardized to their respective untreated cells.

### Size Exclusion Chromatography (SEC) of RecA and EppR

1 mg each of previously purified His-tagged *A. baumannii* RecA (Ching et al., 2024) and EppR (Nguyen et al., 2025) were mixed and dialyzed in SEC running buffer (50 mM HEPES, 300 mM NaCl, 10% glycerol, 2 mM 2-mercaptoethanol, pH 7.5) for 24 h using a 10K MWCO Slide-A-Lyzer^TM^ G3 dialysis cassette (Thermo Scientific). The sample was then concentrated using a 3000 MWCO Vivaspin® 500 centrifugal concentrator (Sartorius) until a final volume of 200 μL. The RecA/EppR mix was loaded onto a Superdex 200 10/300 GL gel filtration column (Separation range: MW 600,000 – 10,000; GE Healthcare Life Sciences) and separated into 1 mL fractions. SEC data was collected and analyzed using UNICORN Control Software (GE Healthcare Life Sciences). Fractions from each defined peak were pooled and concentrated to a final volume of 100 μL using an Amicon® 4 Ultracel-10K centrifugal filter (Millipore). Protein content from each peak was characterized by western blot using anti-RecA and anti- EppR antibodies as described above.

### Growth Curve Measurements

Strains were diluted 1:1000 into Miller-LB medium and grown at 37°C, with shaking at 225 RPM. Colony forming units (CFUs)/mL were measured by spot plating every hour. Measurements were performed in both technical and biological triplicate.

### Electrophoretic Mobility Shift Assay (EMSA)

The *recA* regulatory region (145 bp) was amplified by PCR amplification with primers labeled with a Cy3 fluorophore on the 5’ end (Table 2). DNA (2.5 nM) and purified EppR (Nguyen et al., 2025) were mixed with EMSA binding buffer (10 mM Tris-HCl [pH 8], 10 mM HEPES, 50 mM KCl, 1 mM EDTA, 0.1 mg/mL of bovine serum albumin). EppR was used at the concentrations indicated in figures. All samples include 100 ng of salmon sperm DNA as a nonspecific binding control. Binding reactions were incubated for 1 h at 37°C in a thermocycler (Eppendorf). Reaction mixtures (20 uL) were loaded into a 5% non-denaturing Tris-borate-EDTA (TBE) polyacrylamide gel. Binding complexes were separated at 150 V for 1h at 4°C. Cy3-labelled DNA-protein complexes were visualized using a Bio-Rad Chemidoc imager (Bio-Rad). EMSA experiments were repeated three times and band intensities were quantified using Fiji (Schindelin et al., 2012). The predicted EppR binding site within the *recA* regulatory region was found by scanning the sequence with the previously found conserved EppR binding motif (Nguyen et al., 2025) using the Multiple Em for Motif Elicitation (MEME) program (Bailey et al., 2009).

## Supplemental Figures

**Figure S1:**
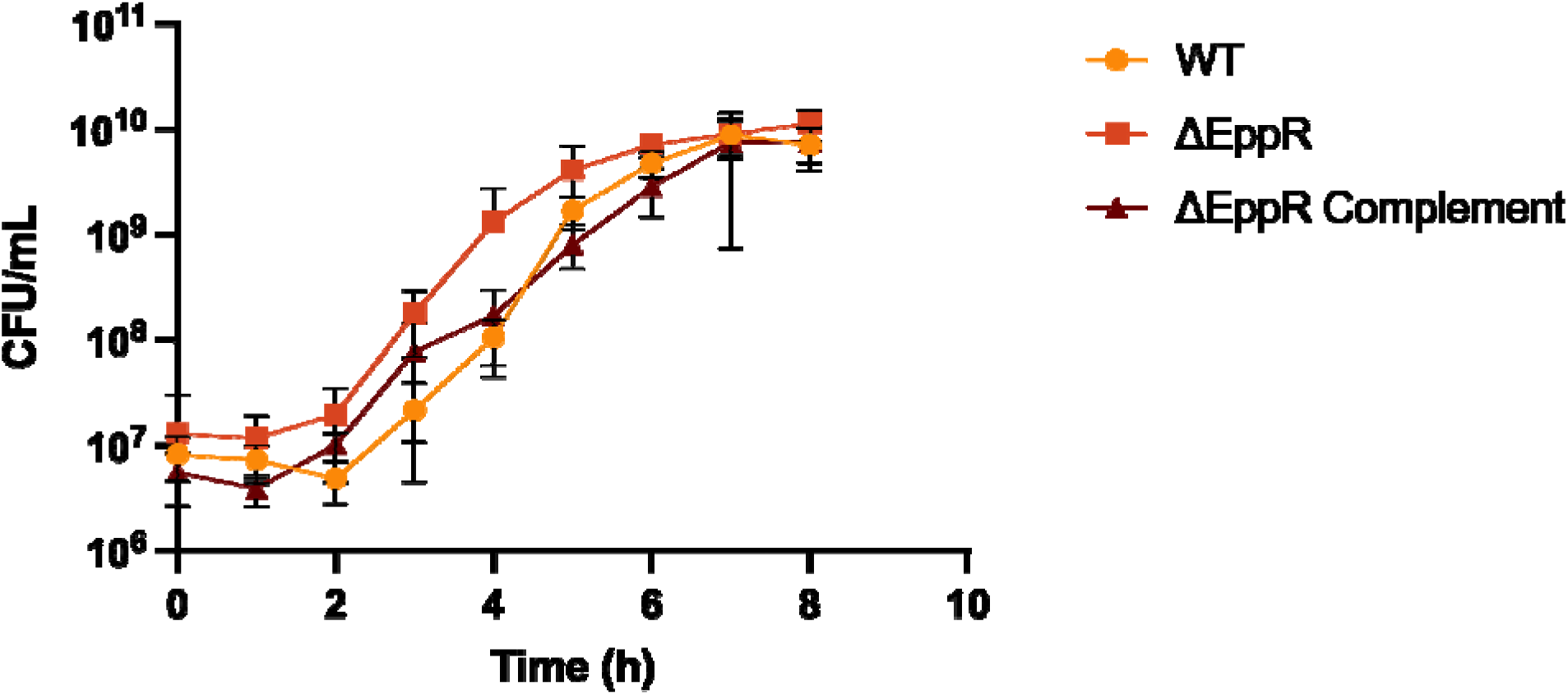
Δ*eppR* does not have a growth defect. Colony forming units (CFUs)/mL was determined as indicated in the Materials and Methods. Error bars represent standard deviation. Measurements were performed in both technical and biological triplicate.

**Figure S2:**
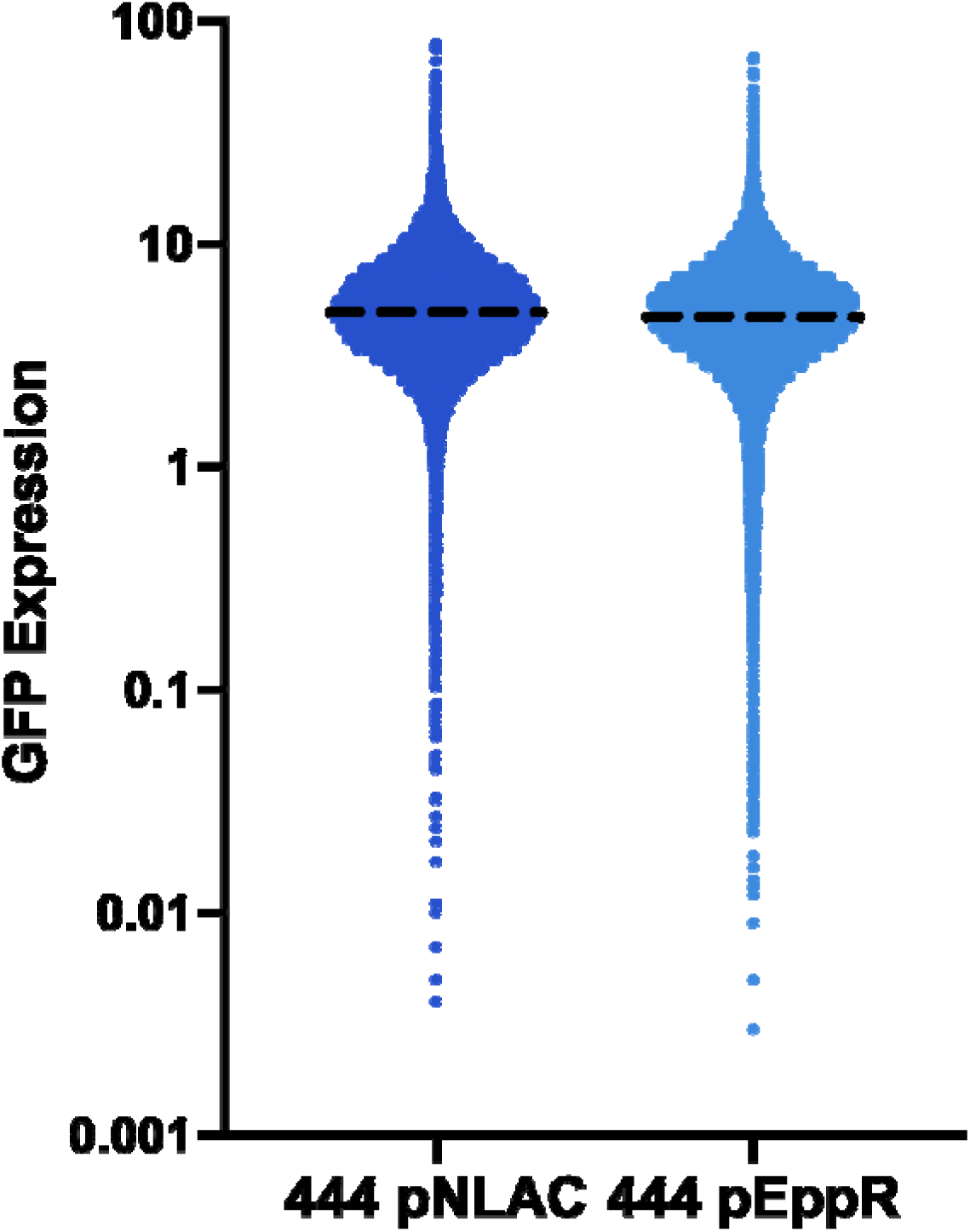
Overproduction of EppR results in a formation of a low RecA expressing subpopulation. Shown here is the single cell *recA* expression from the graph shown in Figure 4B. 444 with pEppR (pNLAC plasmid containing *eppR* under its own promoter) shows a greater fraction of cells that have very low or no fluorescence from a RecA-GFP chromosomal fusion compared to 444 with empty pNLAC. Average expression is denoted by the black dotted line. At least 10,000 cells from 9 independent experiments were quantified using Fiji.

**Figure S3:**
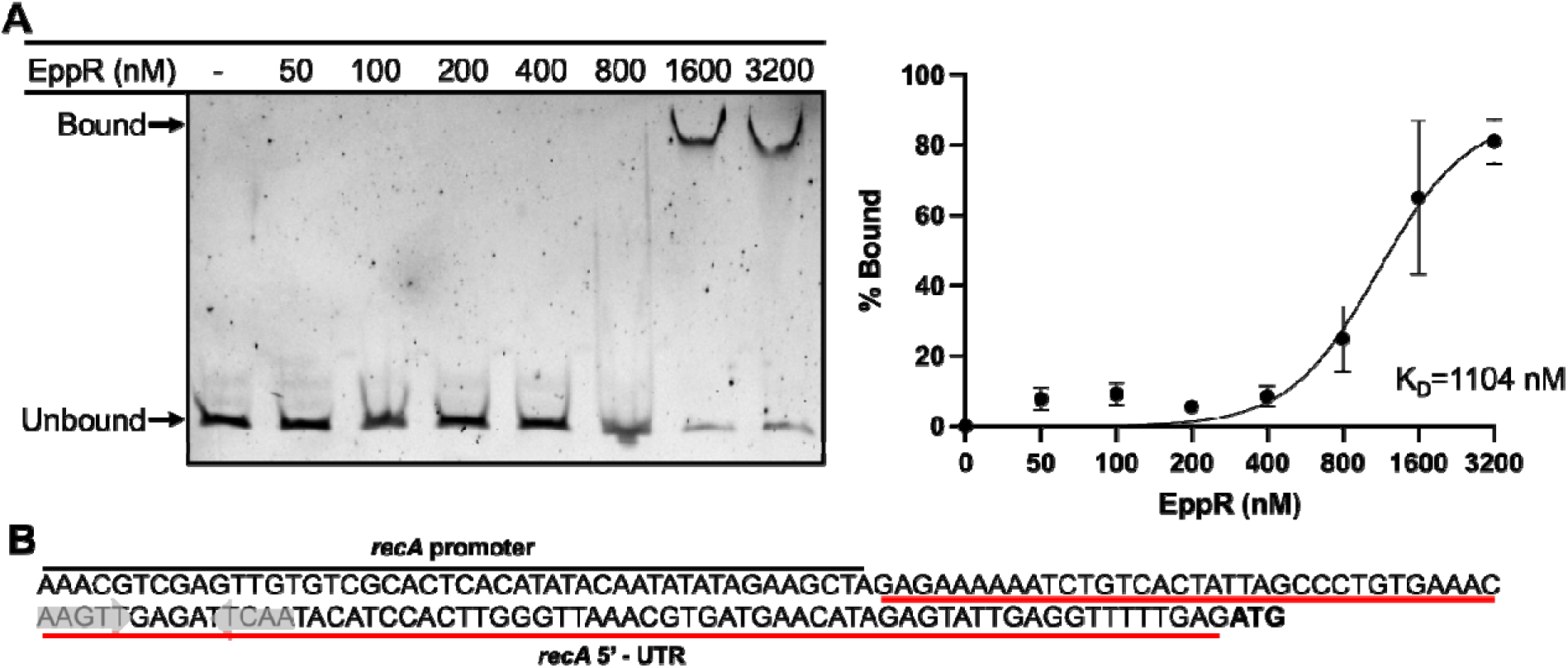
EppR binds to the *recA* regulatory region. (A) EMSA using the *recA* regulatory region shows EppR binding. EMSA experiments were performed with increasing EppR concentrations (0-3200 nM) shown on top of the respective lanes. Gel shown is a representative image of three independent experiments. EMSA was performed as indicated in Material and Methods. Bands representing bound and unbound fractions are denoted on the gel. Full binding was observed at 1600 nM. On the right, quantification of EppR binding to the *recA* regulatory region is shown. Quantification was performed using Fiji and K_D_ was calculated using GraphPad Prism by applying an association kinetics model. Error bars represent standard deviation. (B) A potential EppR binding site was identified in the *recA* regulatory region. The potential binding site was identified using the Motif for Em Elicitation (MEME) program (Bailey et al., 2009) by scanning the *recA* regulatory region with the conserved EppR binding motif that was previously found (Nguyen et al., 2025). The *recA* regulatory region consists of the *recA* promoter and a 5’ untranslated region important for *recA* regulation (Ching et al., 2017). The potential binding site is denoted by gray arrows.

**Figure S4:**
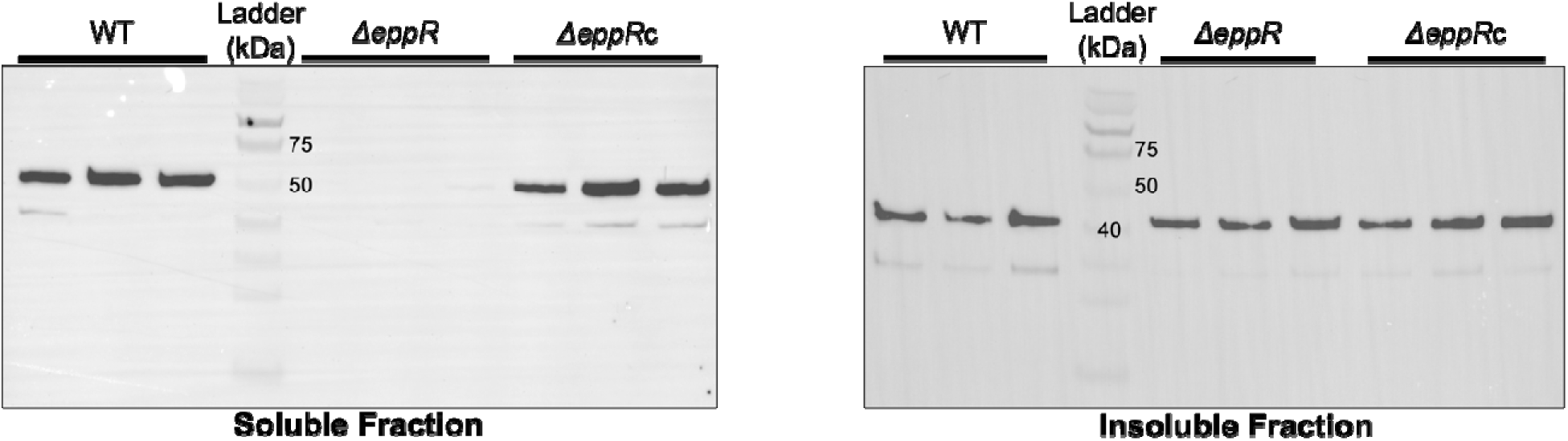
Full images of the RecA western blots shown in. Figure 5A

**Figure S5:**
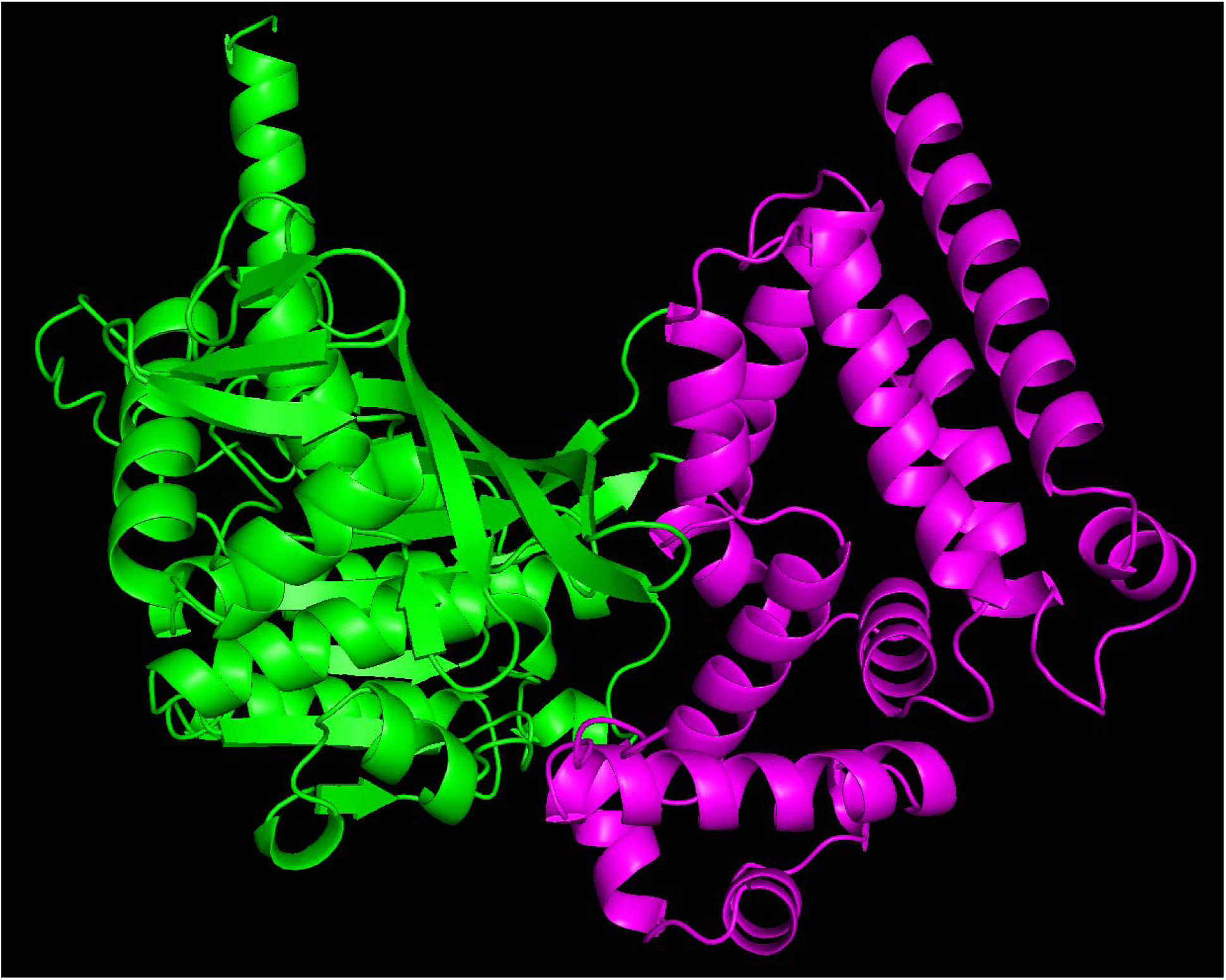
Model of the RecA/EppR complex. Modelling was performed by Alphafold 3. RecA is in green and EppR is in magenta.

## Abbreviations

DDR: **<u>D</u>**NA **<u>d</u>**amage **<u>r</u>**esponse
EppR: **<u>E</u>**rror-**<u>p</u>**rone **<u>p</u>**olymerase **<u>r</u>**egulator
EMSA: **<u>E</u>**lectrophoretic **<u>m</u>**obility **<u>s</u>**hift **<u>a</u>**ssay
SEC: **<u>S</u>**ize **<u>e</u>**xclusion **<u>c</u>**hromatography
WT: **<u>W</u>**ild-**t**ype; *Acinetobacter baumannii* ATCC 17978

## Acknowledgements

V.G-C was funded by a stipend from Northeastern University Skills for Capacity and Inclusion, an Inclusive Excellence award from HHMI.

ALH was funded by the PEAK Summit Award from the Northeastern University Undergraduate Research and Fellowships Office.

We would like to thank the Chai and Geisinger Lab at Northeastern University for their support, feedback, and equipment; Merlin Brychcy for his assistance in editing this manuscript; and the Feldman Lab at Washington University in St. Louis for providing us with the αCsuAB antibody.

